# Nelfinavir inhibits replication of severe acute respiratory syndrome coronavirus 2 in vitro

**DOI:** 10.1101/2020.04.06.026476

**Authors:** Norio Yamamoto, Shutoku Matsuyama, Tyuji Hoshino, Naoki Yamamoto

**Author notes:** Corresponding author: Norio Yamamoto, Department of Virology, Division of Host Defense Mechanism, Tokai University School of Medicine, 143 Shimokasuya, Isehara, Kanagawa 259-1193, Japan, E-mail address or.

## Abstract

In December 2019, severe acute respiratory syndrome coronavirus 2 (SARS-CoV-2) emerged in Wuhan, Hubei Province, China. No specific treatment has been established against coronavirus disease-2019 (COVID-19) so far. Therefore, it is urgently needed to identify effective antiviral agents for the treatment of this disease, and several approved drugs such as lopinavir have been evaluated. Here, we report that nelfinavir, an HIV-1 protease inhibitor, potently inhibits replication of SARS-CoV-2. The effective concentrations for 50% and 90% inhibition (EC50 and EC90) of nelfinavir were 1.13 µM and 1.76 µM respectively, the lowest of the nine HIV-1 protease inhibitors including lopinavir. The trough and peak serum concentrations of nelfinavir were three to six times higher than EC50 of this drug. These results suggest that nelfinavir is a potential candidate drug for the treatment of COVID-19 and should be assessed in patients with COVID-19.

## Introduction

In December 2019, a novel betacoronavirus, designated as severe acute respiratory syndrome coronavirus 2 (SARS-CoV-2), emerged in Wuhan, Hubei Province, China^1,2^. As of March 16, 2020, 167,515 confirmed cases with 6,606 deaths were reported, and 150 countries/territories/areas were affected.

As of now, no specific treatment has been established against coronavirus disease-2019 (COVID-19). After the outbreak of severe acute respiratory syndrome (SARS) in 2003, screening of approved drugs identified at least two human immunodeficiency virus type-1 (HIV-1) protease inhibitors, lopinavir and nelfinavir, as compounds that inhibited SARS-CoV replication in vitro^3,4^. In the open-label study with SARS patients in 2004, the group treated with lopinavir-ritonavir (400 mg and 100 mg, respectively) in addition to ribavirin showed better clinical outcomes than the control group treated with only ribavirin^3^. Because SARS-CoV-2 is relatively similar to SARS-CoV^5^, it is expected that these drugs will be effective for the treatment of COVID-19 patients. To select the candidate drug for the clinical evaluation, the data of antiviral activity in vitro are of great help. However, there is no basic information about the effective concentration of HIV-1 protease inhibitors to abrogate SARS-CoV-2 replication. Thus, in the present study, we evaluated the antiviral activity of nine approved HIV-1 protease inhibitors against SARS-CoV-2 in vitro.

## Results and Discussion

For the evaluation of the nine HIV-1 protease inhibitors, the 50% effective concentration (EC_50_), the 50% cytotoxic concentration (CC_50_), and the selectivity index (SI, CC_50_/EC_50_) were analyzed. Moreover, based on the mean peak and trough serum concentration (C_max_ and C_trough_) of the compounds in human, C_max_-EC_50_ ratio (C_max_/EC_50_) and C_trough_-EC_50_ ratio (C_trough_/EC_50_) were calculated to assess the safety and efficacy of these compounds.

Among these inhibitors tested, the high concentrations of drugs were required to inhibit SARS-CoV-2 replication in amprenavir (EC_50_ = 31.32 µM, CC_50_ > 81 µM, SI > 2.59), darunavir (EC_50_ = 46.41 µM, CC_50_ > 81 µM, SI > 1.75), and indinavir (EC_50_ = 59.14 µM CC_50_ > 81 µM, SI > 1.37) (Fig. 1g, h, i, j). Tipranavir inhibited replication of SARS-CoV-2, but SI was relatively low (EC_50_ = 13.34 µM, CC_50_ = 76.80 µM, SI = 5.76) (Fig. 1f, j). Ritonavir (EC_50_ = 8.63 µM, CC_50_ = 74.11 µM, SI = 8.59), saquinavir (EC_50_ = 8.83 µM, CC_50_ = 44.43 µM, SI = 5.03), and atazanavir (EC_50_ = 9.36 µM, CC_50_ > 81 µM, SI > 8.65) suppressed SARS-CoV-2 at less than 10 µM (Fig. 1c, d, e, j). Lopinavir, which has been clinically tested in patients with SARS and COVID-19, blocked SARS-CoV-2 replication at a low concentration range and its SI was relatively high among nine inhibitors (EC_50_ = 5.73 µM, CC_50_ = 74.44 µM, SI = 12.99) (Fig. 1b, j). Notably, nelfinavir (EC_50_ = 1.13 µM, CC_50_ = 24.32 µM, SI = 21.52) potently inhibited virus replication at the lowest concentration and exhibited the highest SI in the tested HIV-1 protease inhibitors (Fig. 1a, j). To examine the target step of HIV-1 protease inhibitors in the virus life cycle, time-of-addition assays were performed. Intra-cellular viral RNA was quantified at 6 hpi with varying timing of drug addition. Nelfinavir and lopinavir suppressed multiplication of viral RNA in the cells even when these drugs were added at 3.5 hpi (Fig. S1). These results suggest that both drugs functioned at the post-entry step.

**Fig. 1.**
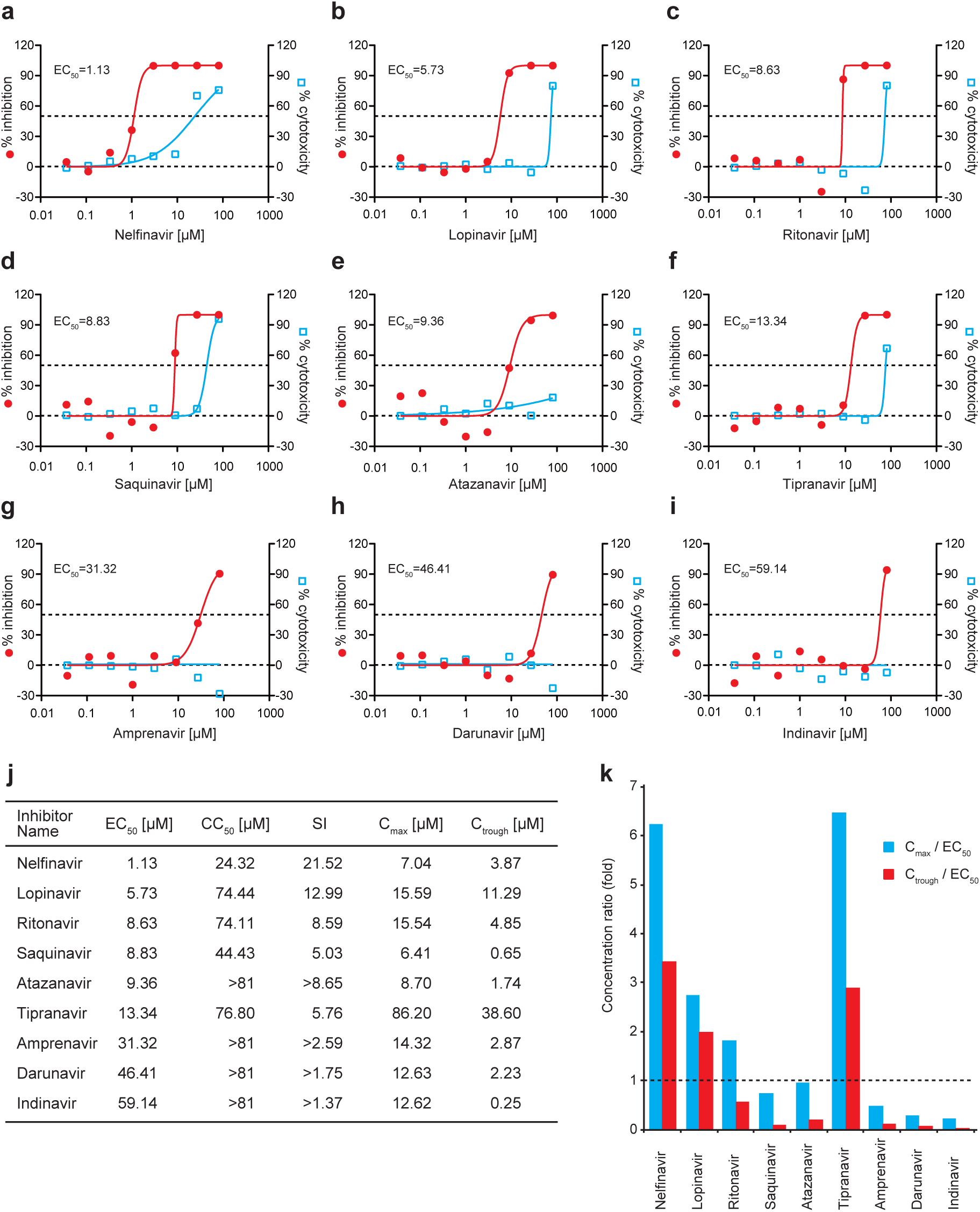
The antiviral activities of HIV-1 protease inhibitors against SARS-CoV-2. **a**-**i**, Vero E6/TMPRSS2 cells were infected with SARS-CoV-2 at a MOI of 0.01 with different concentrations of the indicated protease inhibitors and incubated for 24 h. The viral yield in the cell supernatant was quantified by qRT-PCR. The cytotoxicity of these drugs to Vero E6/TMPRSS2 cells was measured by MTS assays. The left and right Y-axes of the graphs stand for mean % inhibition of the virus yield and mean % cytotoxicity of the drugs, respectively. The experiments were performed in triplicate. **j**, EC_50_, CC_50_, SI, C_max_, and C_trough_ of the HIV-1 protease inhibitors. **k**, C_max_-EC_50_ ratio and C_trough_-EC_50_ ratio of the HIV-1 protease inhibitors.

Next, C_max_-EC_50_ ratio and C_trough_-EC_50_ ratio of nine compounds were calculated to estimate the clinical efficacy against COVID-19. Nelfinavir (C_max_/EC_50_ = 6.23, C_trough_/EC_50_ = 3.43), lopinavir (C_max_/EC_50_ = 2.72, C_trough_/EC_50_ = 1.97), and tipranavir (C_max_/EC_50_ = 6.46, C_trough_/EC_50_ = 2.89) exhibited C_trough_-EC_50_ ratio higher than one (Fig. 1k). Ritonavir exceeded one in C_max_-EC_50_ ratio, but did not in C_trough_-EC_50_ ratio (C_max_/EC_50_ = 1.80, C_trough_/EC_50_ = 0.56). Saquinavir (C_max_/EC_50_ = 0.73, C_trough_/EC_50_ = 0.07), atazanavir (C_max_/EC_50_ = 0.93, C_trough_/EC_50_ = 0.19), amprenavir (C_max_/EC_50_ = 0.46, C_trough_/EC_50_ = 0.09), darunavir (C_max_/EC_50_ = 0.27, C_trough_/EC_50_ = 0.05), and indinavir (C_max_/EC_50_ = 0.21, C_trough_/EC_50_ = 0.004) could not reach one in the C_max_-EC_50_ ratio. These results suggest that nelfinavir, lopinavir, and tipranavir can achieve EC_50_ of each drug in human and are effective in the treatment of COVID-19 patients.

Nelfinavir showed the lowest EC_50_ (1.13 µM), the highest SI (23.32), the second highest C_max_-EC_50_ ratio (6.23), and the highest C_trough_-EC_50_ ratio (3.43) among the tested HIV-1 protease inhibitors. These data indicate that nelfinavir is the best of nine drugs as a potential drug candidate for the treatment of COVID-19. Lopinavir showed the inferior results to nelfinavir, namely, higher EC_50_ (5.73 µM), lower SI (12.99), lower C_max_-EC_50_ ratio (2.72), and lower C_trough_-EC_50_ ratio (1.97) than nelfinavir. In a recent randomized, controlled, open-label trial, lopinavir-ritonavir did not significantly accelerate clinical improvement, reduce 28-day mortality, or decrease throat viral RNA amount in patients with severe COVID-19^6^. One of the reasons why the efficacy of lopinavir-ritonavir was not confirmed in the clinical trial might be associated with insufficient C_max_-EC_50_ and C_trough_-EC_50_ ratio to treat severe COVID-19. It remains to be determined whether earlier lopinavir–ritonavir treatment in mild COVID-19 could have clinical benefit. Tipranavir showed the highest C_max_-EC_50_ ratio (6.46) and the second highest C_trough_-EC_50_ ratio (2.89) of all tested drugs. This is because C_max_ and C_trough_ are much higher than the other tested drugs (Fig. 1j). Since SI of tipranavir was less than that of nelfinavir (5.76 vs. 21.52), nelfinavir seems better than tipranavir. The low SI of tipranavir may raise concern about adverse effects to COVID-19 patients.

Our findings reveal that nelfinavir is highly effective in inhibiting SARS-CoV-2 replication in vitro and has the high C_max_-EC_50_ and C_tough_-EC_50_ ratio. We suggest that nelfinavir is a potential candidate drug for the treatment of COVID-19 and should be assessed in the patients with COVID-19.

## Materials and Methods

### Cells, virus, and antiviral compounds

VeroE6 cells expressing TMPRSS2 (VeroE6/TMPRSS2)^7^ were maintained in Dulbecco’s modified Eagle medium (DMEM; Sigma-Aldrich, St. Louis, MO, USA) supplemented with 5% fetal bovine serum (Thermo Fisher Scientific, Waltham, MA, USA) at 37 °C in a humidified atmosphere of 5% CO_2_. SARS-CoV-2 was propagated in VeroE6/TMPRSS2 cells. All the infection experiments were performed in a biosafety level-3 (BSL-3) laboratory. Nelfinavir a kind gift from Torii pharmaceutical Co. Ltd., and lopinavir was a kind gift from Abbott laboratories. Ritonavir (cat. no. HY-90001), saquinavir (HY-17003), atazanavir (HY-17367A), tipranavir (HY-15148), amprenavir (HY-17430), darunavir (HY-17040), and indinavir (HY-B0689A) were purchased from MedchemExpress (Monmouth Junction, NJ, USA). Camostat (3193, Tocris Bioscience, Minneapolis, MN, USA) and E64d (330005, Calbiochem, San Diego, CA, USA) were used as the entry inhibitors of SARS-CoV-2.

### Evaluation of cytotoxicity and antiviral activity of the drugs

Confluent cells in 96-well plates were treated with the HIV-1 protease inhibitors. After incubation for 24 h, MTS assays were performed to measure the cell viability using CellTiter 96 AQueous One reagent (G3580, Promega, Madison, WI, USA) according to the manufacturer’s instructions. To evaluate the antiviral efficacy of these drugs, confluent cells in 96-well plates were infected with virus at a MOI of 0.01 in the presence of the drugs. After 2 h of incubation, the media with the virus-drug mixture were removed and cells were further cultured with the drug-containing media. The cell culture supernatants collected at 24 hpi was diluted 10-fold in water, then boiled. For time-of-addition assay, drugs were added to cell cultures at 0 or 3.5 hpi, and subsequently cellular RNA was extracted at 6 hpi using a CellAmp Direct RNA Prep Kit (3732, Takara Bio Inc., Shiga, Japan). The qRT-PCR assays were performed to quantify the amount of SARS-CoV-2 RNA with QuantStudio 3 instrument (Thermo Fisher Scientific, Waltham, MA, USA) using primers and probes described previously^8,9^ (N2 set or E set).

## Acknowledgements

We thank Yoko Yamamoto for her technical assistance. This work was supported, in part, by a grant from the Japan Agency for Medical Research and Development to N.Y. [grant no. JP19fk0108051]; and a grant from JSPS KAKENHI [grant no. JP19145824].

## Author contributions

Norio Y. and Naoki Y. conceived and designed the experiments. S.M., Norio Y., and T.H. participated in experiments. Norio Y. and S.M. analyzed the data. Norio Y. wrote the manuscript. Norio Y., S.M., T.H., and Naoki Y. provided the final approval of the manuscript.

## References

1. Huang, C., et al. Clinical features of patients infected with 2019 novel coronavirus in Wuhan, China. Lancet 395, 497–506 (2020).

2. Zhu, N., et al. A Novel Coronavirus from Patients with Pneumonia in China, 2019. N Engl J Med 382, 727–733 (2020).

3. Chu, C.M., et al. Role of lopinavir/ritonavir in the treatment of SARS: initial virological and clinical findings. Thorax 59, 252–256 (2004).

4. Yamamoto, N., et al. HIV protease inhibitor nelfinavir inhibits replication of SARS-associated coronavirus. Biochem Biophys Res Commun 318, 719–725 (2004).

5. Lu, R., et al. Genomic characterisation and epidemiology of 2019 novel coronavirus: implications for virus origins and receptor binding. Lancet 395, 565–574 (2020).

6. Cao, B., et al. A Trial of Lopinavir-Ritonavir in Adults Hospitalized with Severe Covid-19. N Engl J Med (2020).

7. Matsuyama, S., et al. Enhanced isolation of SARS-CoV-2 by TMPRSS2-expressing cells. Proc Natl Acad Sci U S A 117, 7001–7003 (2020).

8. Shirato, K., et al. Development of Genetic Diagnostic Methods for Novel Coronavirus 2019 (nCoV-2019) in Japan. Jpn J Infect Dis (2020).

9. Corman, V.M., et al. Detection of 2019 novel coronavirus (2019-nCoV) by real-time RT-PCR. Euro Surveill 25(2020).

